# An Allosteric Cholesterol Site in Glycine Receptors Characterized Through Molecular Simulations

**DOI:** 10.1101/2024.03.05.583408

**Authors:** Farzaneh Jalalypour, Rebecca J. Howard, Erik Lindahl

## Abstract

Glycine receptors are pentameric ligand-gated ion channels that conduct chloride ions across postsynaptic membranes to facilitate fast inhibitory neurotransmission. In addition to gating by the glycine agonist, interactions with lipids and other compounds in the surrounding membrane environment modulate their function, but molecular details of these interactions remain unclear – in particular for cholesterol. To identify such interactions, here we report on coarse-grained simulations in a model neuronal membrane for three zebrafish glycine-receptor structures, representing apparent resting, open, and desensitized states. We then converted the systems to all-atom models to examine detailed lipid interactions, and observe cholesterol bound to the receptor at an outer-leaflet intersubunit site in a state-dependent manner, indicating that it can bias receptor function. Finally, using a modified perturbation-response scanning approach, we applied short atomistic simulations to identify amino-acid translations correlated with gating conformational changes. Frequent cholesterol contacts in atomistic simulations clustered with residues identified by perturbation analysis and overlapped with mutations influencing channel function and pathology. Cholesterol binding at this site was also observed in a recently reported pig heteromeric glycine receptor. These results indicate state-dependent lipid interactions relevant to allosteric transitions of heteromeric glycine receptors, including specific amino-acid contacts applicable to biophysical modeling and pharmaceutical design.

## Introduction

Pentameric ligand-gated ion channels (pLGICs) mediate cellular communications by converting chemical signals to electrical signals. The classical example involves a presynaptic neuron releasing a neurotransmitter such as acetylcholine or γ-aminobutyric acid, which binds to a pLGIC on the postsynaptic neuron. This agonist interaction induces an allosteric conformational change in the receptor, which leads to pore opening, ion conduction, and membrane potential alteration. ^1^ Apart from their primary agonists, pLGICs are sensitive to range of allosteric modulators, small molecules that can either amplify or reduce the effect of the main agonist.^2–5^ This property has enabled recreational drugs throughout the history of mankind, and it is the basis of several modern clinical drugs including anesthetics.^6–8^ However, the precise mechanisms underlying pLGIC modulation remain unclear.

Members of the pLGIC family can be divided into two main groups: cationic channels, such as the nicotinic acetylcholine receptor (nAChR), which play primarily excitatory roles in neurotransmission; and anionic channels, such as the type-A γ-aminobutyric acid-A receptor (GABAAR)^1^, which feature heavily in neuro-inhibition. Both groups exhibit conserved structural and functional features, including an extracellular domain (ECD) responsible for agonist binding, a transmembrane domain (TMD) forming the central pore for ion conductance, and an intracellular domain (ICD) of variable sequence (Figure 1a). Each TMD subunit contains four helices (M1–M4) connected by extracellular and intracellular loops, and five such subunits create the complete TMD. In the resting state, the channel is closed via a hydrophobic constriction near the middle of the membrane plane. Agonist binding in the ECD triggers a series of conformational changes, including expansion of the hydrophobic gate nearly 50 Å from the neurotransmitter sites, which eventually leads to pore opening. In most eukaryotic pLGICs, this transient open state undergoes a rapid, subtle transition to a desensitized state with the ligand still bound, but where the pore is occluded at an alternative gate facing the cytoplasm. Both gating and allosteric modulation involve ligand binding and conformational changes between and within subunits. ^9,10^.

**Figure 1.**
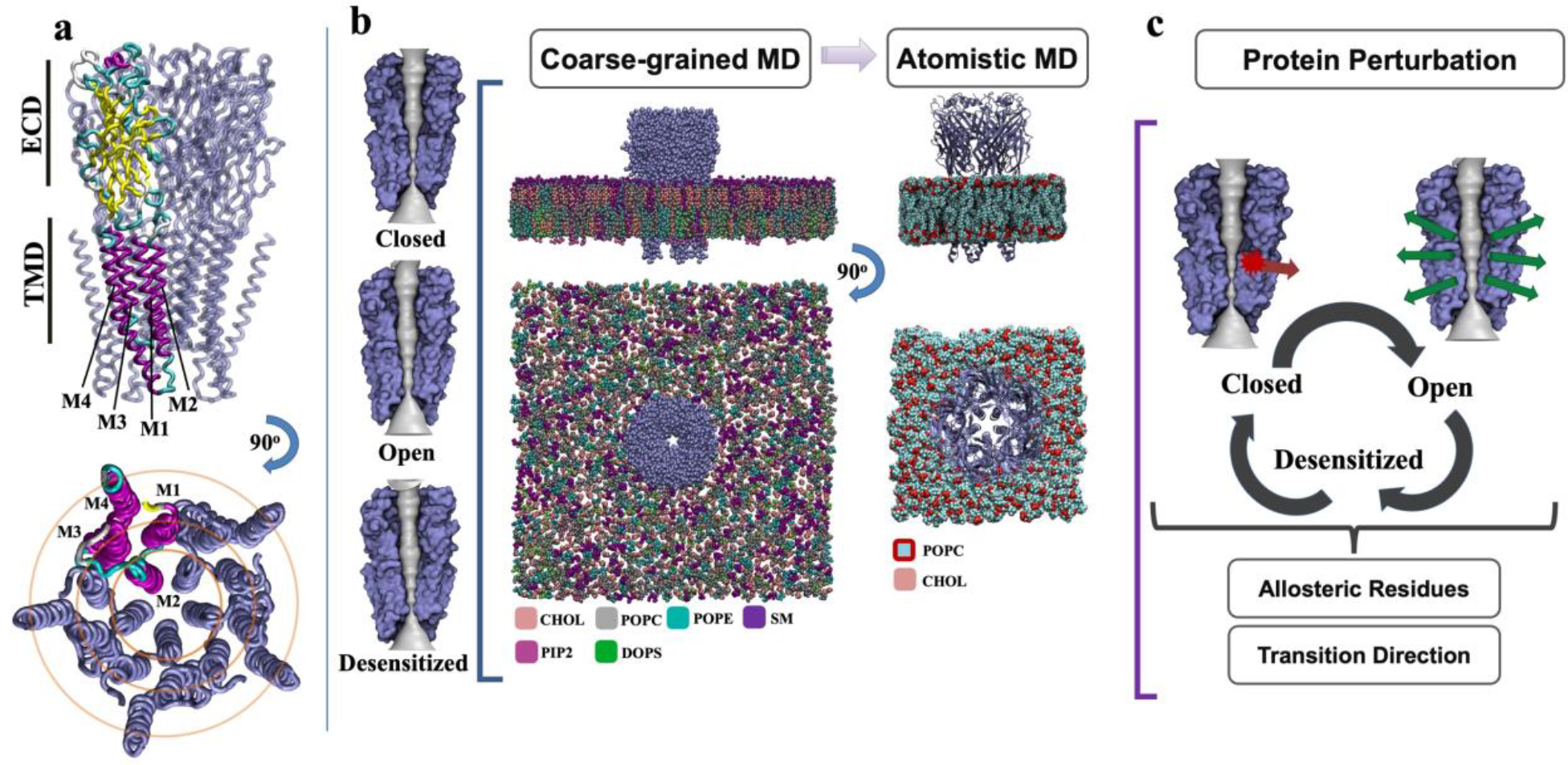
GlyR structural templates and simulation methods. a) Schematic model of a homomeric GlyR (blue), with one subunit colored by secondary structure (yellow, β strands; magenta, ɑ helices; cyan, loops) and transmembrane helices labelled (M1–M4). Due to a lack of structural information, the ICD is not shown. Upper view is from the membrane plane, lower view is of the TMD from the extracellular side. **b)** Molecular simulations of GlyRs in three presumed functional states. Models on the left correspond to GlyR cryo-EM structures reported in closed (PDB ID 6PM3), open (PDB ID 6PM2), and desensitized (PDB ID 6PM1) states, all in the presence of taurine. Structures are represented as surfaces (blue), with the foremost two subunits removed for clarity, revealing the linear pore (gray). Center views are of a representative coarse-grained simulation box, with multiatom beads representing the receptor (blue) and six different types of lipids (colored separately according to legend). The rightmost view illustrates an atomistic simulation of a back- mapped conformation, with the receptor represented as ribbons (blue) and cholesterol (pink) as well as POPC (cyan, colored by heteroatom) as spheres. Water and ions are removed for clarity. **c)** Schematic of protein perturbation calculations, in which a perturbation (red arrow) induces motions of residues (green arrows) that are assessed by their overlap with the direction of a relevant conformational change, in this case between closed, open, and desensitized states.

Recent structures of the glycine receptor (GlyR), an anionic pLGIC, provide notable insights into the architecture and functional cycling of these channels. The GlyR is a major inhibitory chloride channel in the spinal cord and brain and it is the target of several allosteric modulators. ^2,3^ Functional GlyRs include both homomeric assemblies of α1, α2, or α3 subunits^11,12^ and heteromeric combinations of four α subunits and one β subunit.^13–15^ One recent study reported structures of the homomeric zebrafish α1 GlyR in multiple apparent functional states, including conformations annotated as closed, open, closed, desensitized, and “super-open,” processed from the same cryo-EM data set in the presence of the partial agonist taurine. ^11^ These receptors were extracted directly from insect cell membranes using styrene maleic acid (SMA) copolymer, enabling the preservation of embedding lipids from the expression conditions, although in this case no specific lipids were resolved in the final structures. Shortly thereafter, a structure of the native pig α1β GlyR was reported in an apparent desensitized state with the agonist glycine ^13^. Subsequent agonist-bound structures of heteromeric GlyRs from zebrafish and pig largely recapitulated this asymmetric desensitized state. ^14,15^ These structures provide opportunities to investigate state-dependent lipid interactions and allostery in both homomeric and heteromeric systems.

Lipid interactions have received increasing attention in structure-function studies of membrane proteins. First, general physical characteristics of the lipid bilayer, such as fluidity, thickness, and curvature, have been shown to influence membrane-protein properties. ^16,17^ For instance, membrane thinning by rhomboid proteins has been shown to modify elastic properties required for rhodopsin function. ^18^ Further, specific lipid types have been shown to bind selectively to transmembrane protein surfaces and modulate function. For example, specific phospholipids allosterically modulate G-protein coupled receptors (GPCRs) ^19^, and polyunsaturated fatty acids have been shown to inhibit pLGICs ^20^ or altering the sensitivity of voltage-gated ion channels^21^. Notably, the steroid lipid cholesterol stabilizes structural elements of the β2 adrenergic receptor, a prototypical GPCR, and hinders sampling of its conformational landscape. ^22^ The sodium–potassium ATPase pump is also more active when cholesterol is present. Function of the nAChR is similarly dependent on lipid content as well as bulk fluidity, particularly on the presence of cholesterol. ^23–25^ For LGICs, cholesterol has been shown to enhance agonist-induced channel opening as well as desensitization of the cationic nAChR, while its depletion reduces function. ^25,26^ Cholesterol has also been proposed to bind specific sites in anionic pLGICs ^25,27,28^, but it has this far been challenging to retain bound lipids during reconstitution of pLGICs for structure determination of complexes.

As an alternative approach, modern simulation methods may offer a range of insights into the complex dynamics of membrane-protein system such as the GlyR. For instance, it has been shown that lipid localization and interactions with proteins can be predicted by long MD simulations. ^16,29^ While coarse-grained simulations in particular remove much of the detailed interactions by grouping several atoms as beads to reduce the degrees of freedom ^30,31^, this approximation enables them to cover significantly longer timescales while retaining at least some qualitative accuracy. This can provide excellent sampling in complex systems limited by diffusion, and the results of coarse-grained simulations can then be converted back to all-atom models and used to initiate atomistic simulations to provide more detailed models of interactions e.g. with lipids in specific sites in a dynamic protein. Coarse-grained simulations starting from homology models of homopentameric human GlyRs have recently been used to indicated cholesterol preferentially binds to the open over the closed state. ^28^

Here, we took advantage of recent GlyR structures in closed, open, and desensitized states determined under identical, nanodisc-embedded conditions to study computational cholesterol-protein dynamics.^11^ We leveraged coarse-grained and atomistic MD simulations (Figure 1b) to identify specific sites occupied by cholesterol in particular functional states throughout the conformational cycle. We also investigated cholesterol interactions with the heteromeric GlyR, and related our findings to previous functional data, particularly in synthetic and disease variants. Since coarse-grained simulations require scaffold restraints on the protein structure and atomistic simulations struggle to cover timescales relevant for gating, it is difficult to directly simulate the complete gating and allosteric modulation transition mechanisms. Thus, to locate potential regions and residues of interest, we chose as a subsequent step to employ a perturbation approach to identify residues whose motions correlate with the conformational cycling. Briefly, this method generates the responses of all residues in a protein to a perturbation inserted at selected site. These perturbations mimic external forces, such as random Brownian kicks or ligand binding. The objective is to find so-called allosteric residues whose displacements overlap most with the conformational change between two states of a protein. This approach has been used to locate hotspots that influence protein dynamics in multiple systems ^32,33^ , and the perturbation analysis (Figure 1c) enabled us to identify specific sites occupied by cholesterol in particular functional states, and amino acid residues likely to allosterically mediate cholesterol effects on conformational change. Our results identify specific state-dependent lipid interactions likely to influencing the allosteric transitions of the GlyR embedded in a biologically realistic membrane, and amino acids contacts that are potential targets for pharmaceutical design.

## Materials and Methods

### Coarse-grained starting models

For simulations of homomeric GlyRs, starting models were taken from structures of SMA-extracted zebrafish α1 GlyRs in the presence of taurine. ^11^ The selected models were classified from the same cryo-EM dataset as closed (PDB ID 6PM3), open (PDB ID 6PM2), and desensitized (PDB ID 6PM1)^11^ (Table 1). Simulations of the heteromeric GlyR were based on the pig α1β GlyR, whose amino acid sequence is 99% identical to the human GlyR, in the presence of glycine, solubilized from native spinal cord in n-dodecyl-β-maltoside, and classified by cryo-EM as desensitized (PDB ID 7MLY). ^13^ Since the ICD was not resolved in any of the experimental structures, it was replaced in our models by a short, flexible loop (Ala-328, Gly-329, Thr-330) ^34^ using MODELLER. ^31^

**Table 1.** Starting models used for simulations in this study.

| PDB ID | Type | Subunit(s) | Structural state | Organism | Resolution (Å) |
| --- | --- | --- | --- | --- | --- |
| **6PM1** | Homopentameric | $\alpha 1$ | Desensitized | *Danio rerio* | 3.0 |
| **6PM2** | Homopentameric | $\alpha 1$ | Open | *Danio rerio* | 3.0 |
| **6PM3** | Homopentameric | $\alpha 1$ | Closed | *Danio rerio* | 3.0 |
| **7MLY** | Heteropentameric | 1 $\beta$/4 $\alpha 1$ | Desensitized | *Sus scrofa* | 2.7 |

### Equilibrium coarse-grained MD simulations in an asymmetrical neuronal membrane

The MARTINI force field (Martini 2.2 amino acid, Martini 2.0 lipids and non-polarizable water) ^31^ was employed for coarse-grained simulations, in which one backbone bead and 0–3 side chain beads represent each residue. The Martini Bilayer Maker in CHARMM-GUI ^35^ was used to insert each protein structure (closed, open, or desensitized) in an asymmetrical bilayer of lipids that resembled the neuronal membrane (Table 2), by using a lipid composition taken from previous studies. ^28,36^ The outer leaflet contained 44.9% cholesterol (CHOL), 24.4% phosphatidylcholine (POPC), 11.2% phosphoethanolamine (POPE), and 19.5% sphingomyelin (SM), while the inner leaflet consisted of 43.3% CHOL, 13.4% POPC, 21.6% POPE, 3.1% SM, 16.5% dioleoylphosphatidylserine (DOPS), and 2.1% phosphatidylinositol 4,5-bisphosphate (PIP2). In all coarse-grained simulations, taurine ligands were removed. The systems were neutralized by addition of ions to approximate 150 mM NaCl. In total, the box spanned 256 × 256 × 169 Å containing protein, lipids, ions, and nonpolarizable water. After energy minimization and equilibration, involving a gradual lowering of the force of the restraints, four replicas of each system were simulated for 22 μs each using GROMACS 2021 ^37^, with CG scaffolds to restrain the protein secondary structure while lipids, ions, and water were freely diffusing.

**Table 2.** Lipid composition of the asymmetrical neuronal membrane mimic.

| **Lipid type** | **Number in outer leaflet (%)** | **Number in inner leaflet (%)** |
| --- | --- | --- |
| **CHOL** | 528 (45%) | 504 (43.3%) |
| **POPC** | 288 (24.4%) | 156 (13.4%) |
| **POPE** | 132 (11.2%) | 252 (21.6%) |
| **DOPS** | 0 (0%) | 192 (16.5%) |
| **PIP2** | 0 (0%) | 24 (2.1%) |
| **SM** | 228 (19.4%) | 36 (3.1%) |
| **Total** | 1176 (100%) | 1164 (100%) |

### Occupancy analysis and binding site identification

Protein-lipid interactions were assessed using two different approaches. First, lipid occupancies during coarse-grained simulations were measured for each structural state. To this end, the distribution of each lipid type was estimated from the four 22-μs replicates and averaged over all frames using the Volmap tool in VMD. ^38^ A grid with a resolution of 1 Å in each dimension was used to calculate lipid occupancy. Lipid density was reported when lipid and protein were in contact in a given position in at least 50% of simulation frames. Second, PyLipID ^39^, a Python-based analysis tool, was used to further examine lipid sites including native-like binding poses and the interaction residence time of each residue.

### Atomistic starting models

To generate atomistic starting poses for each structure with cholesterol in its putative interaction site, we first back-mapped the final frame of an open-state coarse-grained simulation, including the protein and its five most closely associated cholesterol molecules, to atomistic representation in POPC using the CG2AA tool ^40^ together with the bilayer builder in CHARMM-GUI. ^35^ Following system equilibration, we then ran a preliminary 300-ns fully unrestrained simulation to relax cholesterol interactions at the protein-lipid interface. To obtain a reasonably symmetric starting state with similar binding in all sites of the homomeric channel, we then selected the cholesterol molecule with the lowest root-mean-squared deviation (RMSD) from its initial pose as a best fit to the interfacial site and inserted it symmetrically at all five subunit interfaces of the closed, open, and desensitized starting models.

### Atomistic MD simulations

To assess cholesterol binding, atomistic simulations of cholesterol-bound receptors in each state (prepared as described above) were simulated in quadruple replicates covering 300 ns each.

Atomistic MD simulations.^35^ Simulations were performed using GROMACS 2021 ^37^ with CHARMM36 force field parameters ^41^, and the TIP3P water model. The systems were neutralized by adding ions to approximate 150 mM NaCl, and he simulation timestep set to 2 fs. The bilayer dimensions were 120 × 120 × 170 Å. LINCS ^42^ was used to constrain the length of hydrogen bonds. The particle-mesh Ewald approach ^43^ was used to estimate long-range electrostatic interactions. The Parrinello–Rahman barostat ^44^ and v-rescale thermostat ^45^ were used to maintain pressure (1 bar) and temperature (300 K), respectively. Pore characteristics in each state were analyzed using the channel annotation package (CHAP). ^46^

### Atomistic contact and interaction assessment

Cholesterol contacts with the receptor were first assessed by measuring the distance between the oxygen atom of the hydroxyl group of each cholesterol molecule and the hydrogen atom (HG1) of residue Ser-283 using Python MDAnalysis scripts.^47^ Then, the Protein-Ligand Interaction Fingerprints (ProLIF 1.0.0) tool ^48^ was used to generate an interaction map, screening all potential interactions using the default distance cutoffs of 3.5 Å for hydrogen bonds and 4.5 Å for hydrophobic, π–π, and cation–π interactions.

### Protein perturbation calculations

In order to investigate protein dynamics, a second set of simulations was initiated from the same atomistic starting structures but removing the glycine agonist and again performing quadruple replicate simulations for 300 ns in each state. The technique of protein perturbation, which has previously been used to investigate conformational modulation of proteins. ^32,33,49,50^ This method relies on a covariance matrix, derived from MD simulations, to relate external forces to shifts in atomic coordinates according to the principles of linear response theory. The coarse-grained representation of each state was constructed by considering each Cα atom as a node. Then, numerous random forces were sequentially applied in various directions to each node, generating displacement values. To attain a desired state, the objective is to identify a residue and the direction in which it needs to be perturbed to generate effective displacements. The anticipated displacements are compared to the difference between closed and open structures, which in turn is used to identify residues whose response to perturbations have high overlap (*O*^*i*^) with the gating conformational transition as candidate hotspots, where previous studies have indicated ∼0.6 as a threshold for significance.^32^

In this study, protein RMSD and pore characteristics of GlyR models were first measured to verify the stability of the structure during simulations (Figure S1). Parameters including trajectory interval, and number of perturbations were optimized based on previous protocols ^32^ to maximize sampling and achieve converged overlap values. Each residue of the pentameric receptor was subsequently perturbed by 1000 different force vectors. Three separate calculations were done to cover the closed-to-open, open-to-desensitized, and desensitized-to-closed conformational transitions. For each transition, an equilibrated structure was chosen from the frames within the simulation to serve as the starting point for the transition, since the perturbation approach requires a minimized and relaxed conformation as described by the force field. The experimental structures were used as a target states. The perturbation approach applies random forces, and sampling was enhanced by repeating each (random force) perturbation calculation five times for each transition. Since the structural difference between the open and desensitized conformations is limited (mostly localized to a ∼1Å contraction in the inner pore radius), and because this transition is not expected to involve communication between ECD and TMD, we do not expect this transition to indicate any significant overlap values, but we rather include it as a reference to check that the perturbation approach does not generate spurious high correlations.

## Results

### A state-dependent site for cholesterol identified by coarse-grained simulations

To characterize lipid interactions in all three functional states, we first applied coarse-grained MD simulations to cryo-EM structures of the lipid-embedded full-length zebrafish α1 GlyR (PDB IDs 6PM3, 6PM2, and 6PM1) determined at ≤3.2 Å resolution from three distinct classes of the same cryo-EM dataset ^11^. Although all the above-mentioned structures contained the partial agonist taurine in the extracellular ligand-binding site, the TMDs were notably distinct from one another in particular between resting and open states. The first was closed, similar to a resting state; the other two were comparable to open and desensitized structures determined in the presence of other partial or full agonists. In the context of lipid interactions, which occur primarily in the TMD, these three structures were therefore presumed to represent closed, open, and desensitized states.

In order to simulate interactions with physiologically relevant lipids, we first embedded each structure in an asymmetric bilayer designed to mimic a neuronal membrane, ^28,36^ and ran four replicate 22-μs simulations of each structure (Figure 1b). Filtering at 50% occupancy, no strongly preferred sites were systematically observed for phospholipids, including POPC, POPE, DOPS, SM, or PIP2. Conversely, a groove buried in the membrane outer leaflet between the M2 and M3 helices of one subunit and the M1 helix of the complementary subunit was distinctively occupied by cholesterol in simulations of both the open and desensitized states (Figure 2a, b). This interaction appeared to be state-dependent; in the closed state, cholesterol was only observed to have weak occupancy around M4.

**Figure 2.**
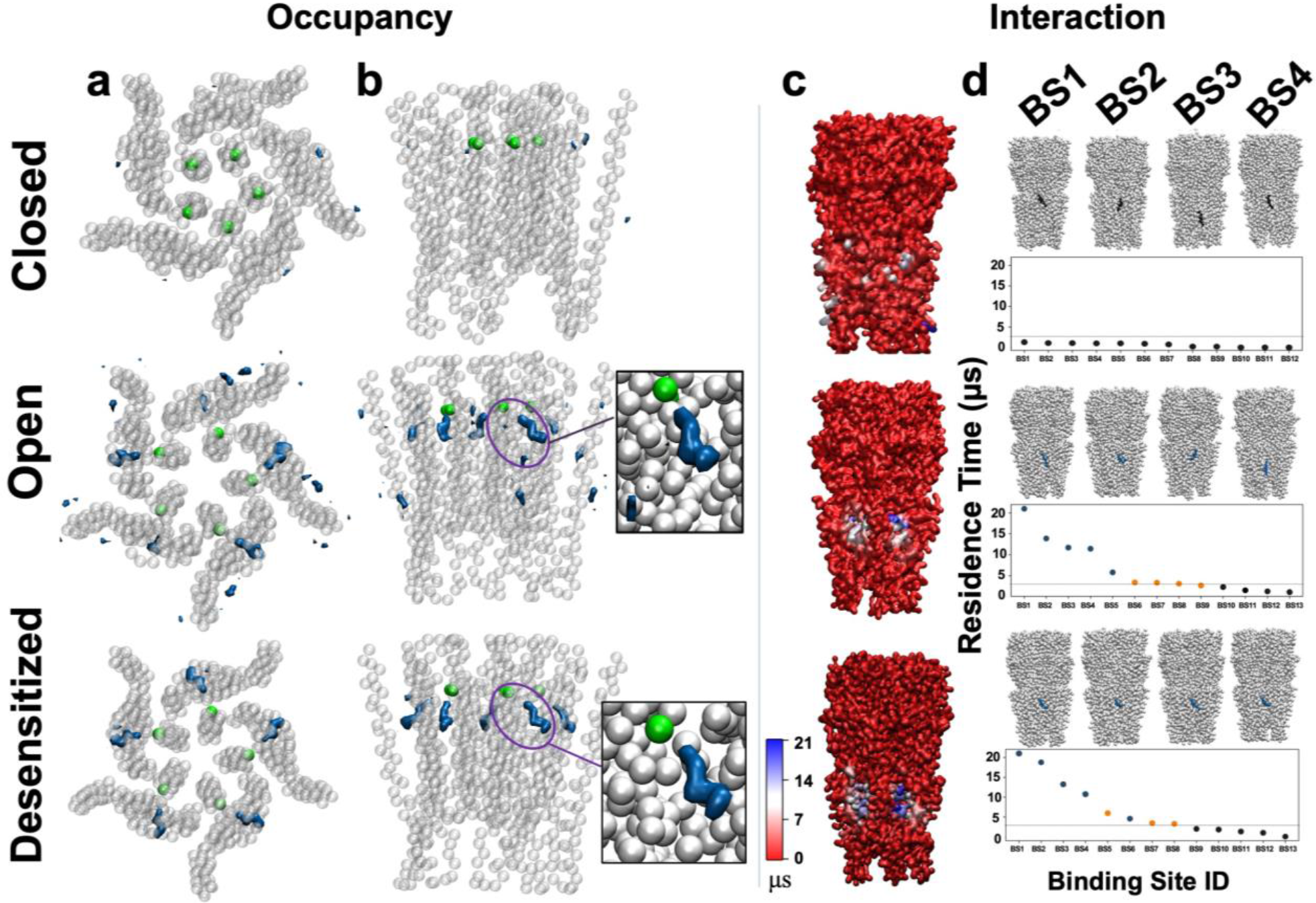
A state-dependent site for cholesterol identified by coarse-grained simulations. a) Densities (blue) representing >50% cholesterol occupancy around the GlyR TMD (gray), including the bead representing Ser-283 (green), viewed from the extracellular side. **b)** Densities as in panel *a*, viewed from the membrane plane. Insets show zoom views of the high-occupancy region in the open and desensitized states. **c)** Interaction residence times of cholesterol with protein residues calculated by PyLipID, shown as a molecular surface colored according to the scale at bottom left (red–blue, 0– 21 μs). **d)** Interaction residence times of cholesterol at the top twelve individual PyLipID-identified binding poses. Poses with the four longest residence times (BS1–BS4) are illustrated above each plot, showing representative conformations of the protein (gray) and cholesterol (blue). Interaction times are shown for the five symmetrical intersubunit sites (cyan), as well as alternative sites of more (brown) or less (gray) than 3 μs (threshold line). Each column panel shows data for the closed (PDB ID 6PM3, top), open (PDB ID 6PM2, middle) and desensitized (PDB ID 6PM1, bottom) states.

To verify this apparent state-dependent cholesterol site and quantify its dynamics, we used the PyLipID tool ^39^ to identify discrete sites associated with the longest protein-interaction residence times across all coarse-grained simulations of each system. Distinctive interaction hotspots for cholesterol were identified in both open and desensitized structures at the outer intersubunit cleft (Figure 2c and Figure S2), corresponding to the regions of high occupancy described above. Indeed, in both states the longest-lived cholesterol interaction poses — each with at least 3 μs residence time — overlapped the high-occupancy site in at least four of the five equivalent subunit interfaces for all simulations (Figure 2d). Conversely, cholesterol interactions in the closed state were far shorter-lived and broadly distributed.

### Cholesterol stability and interactions revealed by atomistic simulations

Having identified a putative cholesterol binding site specific to open and desensitized states of the receptor, we next sought to characterize amino acid contacts with cholesterol in detail using atomistic simulations in the smaller POPC bilayer (see Methods). In the initial cholesterol pose, the hydroxyl moiety of cholesterol was oriented toward Ser-283 on the M2 helix of each subunit, with its multi-ring system bridging the subunit cleft and its hydrophobic tail projecting towards the lipid bilayer (Figure 3a). During the quadruplicate 300 ns simulations, cholesterol remained relatively close to Ser-283 in simulations of both the open and desensitized states (Figure 3b–c). Conversely, cholesterol was rapidly displaced from this initial orientation in closed-state simulations, often flipping, reorienting, or dissociating entirely from the receptor.

**Figure 3.**
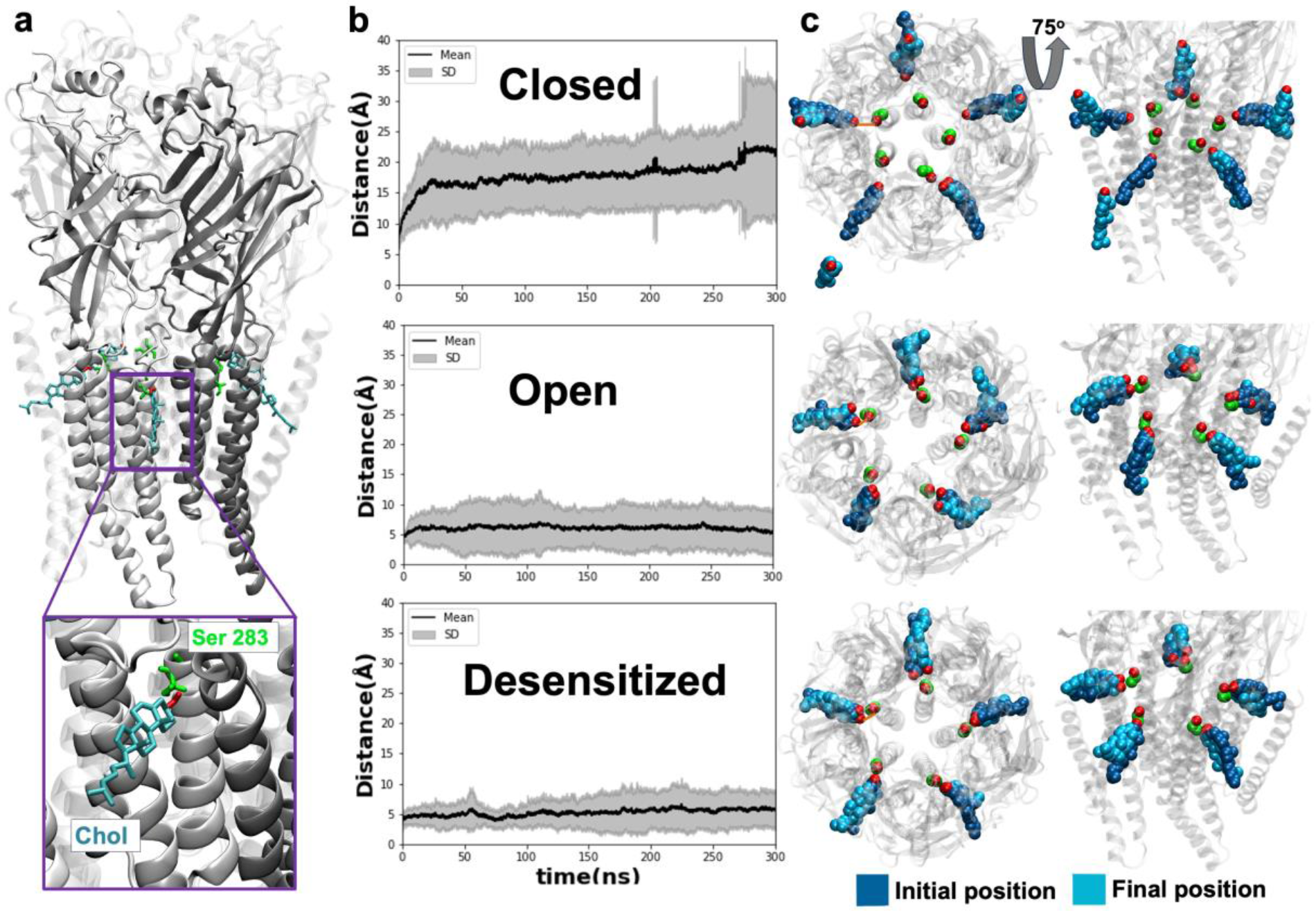
Cholesterol stability and interactions revealed by atomistic simulations. a) Representative all-atom starting model of the GlyR (gray) with cholesterol molecules (Chol, cyan) bound at the intersubunit sites, proximal to Ser-283 (green). Inset shows a zoom view of a single intersubunit site. For clarity, the principal and complementary subunits in the zoom-view site are shown as light and dark opaque ribbons, with the remaining subunit semi-transparent. Cholesterol and Ser-283 are shown as licorice, colored by heteroatom. **b)** Mean distance (± standard deviation, SD, gray) between Ser-283 and its adjacent cholesterol molecule for all five subunits in all four replicas in each state. **c)** Initial (1 ns, blue) and final (300 ns, cyan) positions of cholesterol molecules, colored by heteroatom, in representative atomistic simulations of GlyRs (gray) in three states. Lefthand views show each system from the extracellular side, righthand views from the membrane plane.

Amino acid interaction fingerprints generated in ProLIF ^48^ identified a discrete cholesterol pocket surrounded by the M2 and M3 helices of one subunit and M1 of the complementary subunit in both open and desensitized states (Figure 4). In particular, residues Ser-283, Arg-287, and Asp-300 were capable of polar interactions with the cholesterol hydroxyl group, while bands of neighboring residues including Ile-241, Ile-245, Pro-246, Leu-249, and Ile-252 on M1; Thr-280 on M2; and Val-296, Ile- 301, Met-303, Ala-304, Val-305, Leu-307, Leu308, Phe-311, and Leu-315 on M3 made frequent hydrophobic contacts with the steroid.

**Figure 4.**
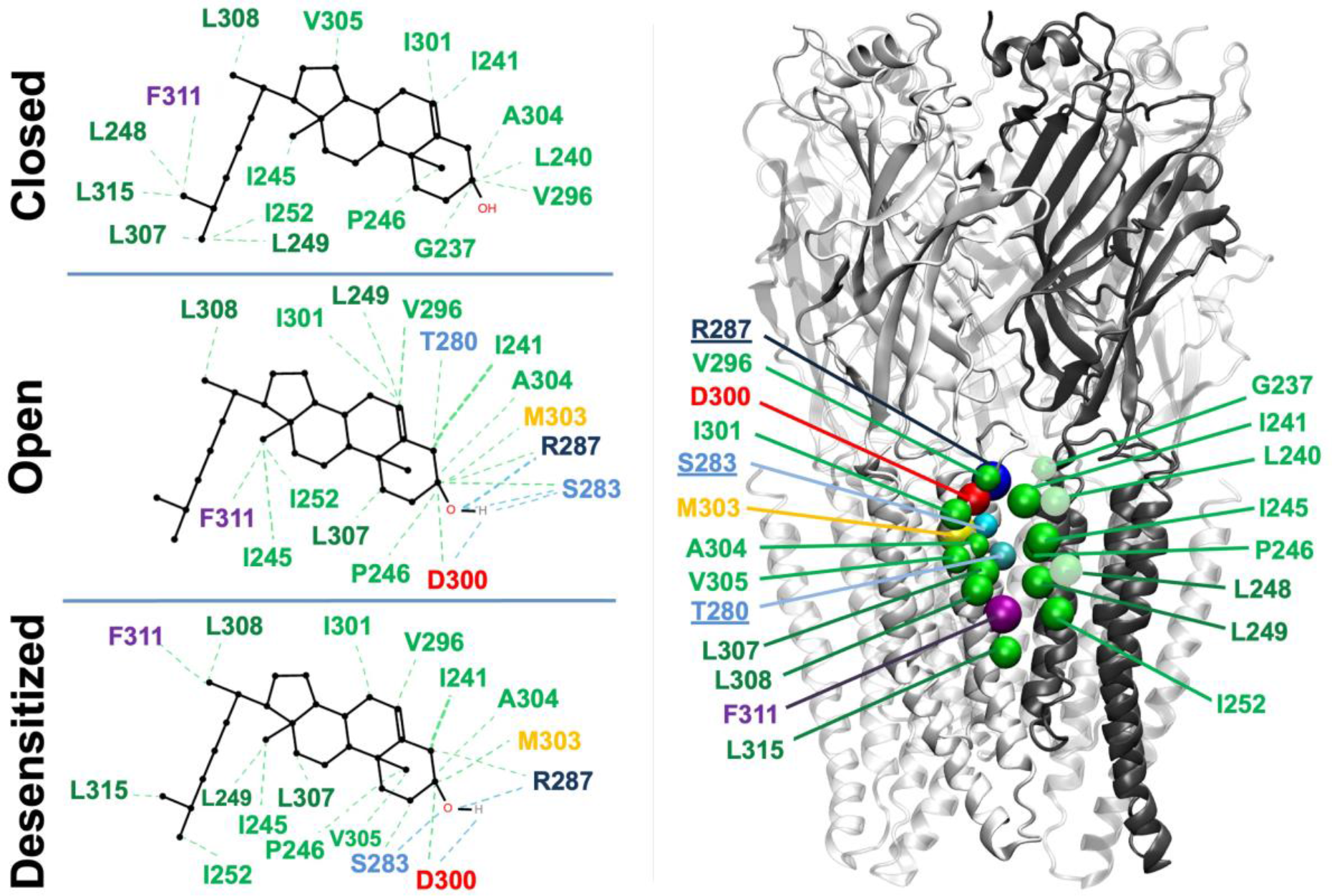
Cholesterol contacts at the outer-leaflet interfacial site. Lefthand panels show interaction fingerprints for cholesterol generated in ProLIF from quadruplicate all-atom MD simulations of closed (top), open (middle), and desensitized (bottom) states of the receptor. Righthand panel shows residues highlighted by ProLIF as Cα spheres on a representative GlyR model, depicted as in *Figure 3a*. Underlined residues are located on M2. The open and desensitized states share similar patterns, whereas the closed state displays weaker interactions and lacks hydrogen bond M2 contacts. In all panels, residues are colored by chemical property, including hydrophobic (green), aromatic (purple), acidic (red), polar (cyan), basic (navy), and sulfur-containing (yellow) side chains.

### Cholesterol-binding region implicated in channel gating by protein perturbation analysis

To independently identify glycine-receptor residues of potential relevance for allosteric gating, we used a modified perturbation-response scanning approach ^32,33^ based on additional atomistic simulations of each functional state without glycine (see Methods). The overall protein did not deviate substantially from its initial structure (within 3.5 Å Cα-RMSD) or alter its pore conformation. The perturbation-scanning approach identified high overlap scores (>0.6) both for the opening ^32^(closed-to-open) and recovery (desensitized-to-closed) transitions (Figure S1, Table 3) involving distinctive residue sets, as detailed below.

**Table 3.** Summary of protein perturbation results including highest overlap values and key residues.

| Protein perturbation | Initial structure ID (simulation frame) | Target structure ID | Converged trajectory interval | Average Cα-RMSD in interval | Best overlap (*O<sub>i</sub>*) | Key residues |
| --- | --- | --- | --- | --- | --- | --- |
| **Closed-to-open** | 6PM3 (180 ns) | 6PM2 | 180–300 ns | 2.8 Å | 0.65 - 0.60 | **305, 308, 414, 255, 251, 409, 410, 306** |
| **Open-to-desensitized\*** | 6PM2 (60 ns) | 6PM1 | 60–300 ns | 3.1 Å | 0.43 - 0.40 | **117, 171, 170, 229, 205, 167** |
| **Desensitized-to-closed** | 6PM1 (150 ns) | 6PM3 | 150–300 ns | 2.8 Å | 0.72 - 0.70 | **277, 278, 242, 237, 238, 163, 240, 267, 244** |

\*The transition between open and desensitized states was not associated with any key residues with overlap >0.40, consistent with this being a relatively subtle/localized conformational change.

For the closed-to-open transition, perturbation analysis identified M1 residues Val-251 and Trp- 255, M3 residues Val-305, Cys-306, and Leu-308, and M4 residues Ala-409, Phe-410 and Phe-414 as potentially allosterically relevant, with an overlap of 0.60–0.65 (Table 3, Figure S3). Notably, Val- 305 and Leu-308 were also frequent direct contacts with cholesterol in atomistic simulations, while Val-251, Trp-255, and Phe-414 occupied a proximal shell around the cholesterol site (Figure 5). For the recovery desensitized-to-closed state transition, relevant residues identified by perturbation analysis included Met-163 in the extracellular Cys loop, Gly-237, Tyr-238, Leu-240, Gln-242, and Tyr-244 in M1, and Ala-267, Leu-277, and Thr-278 in M2, with an overlap of 0.70–0.72 (Table 3, Figure S3). Again, several of these residues clustered around the proposed state-dependent cholesterol site (Figure 5). Force vectors associated with the highest-overlap residues relevant to both opening and recovery transitions were near perpendicular to the pore axis, implying a blooming transition at the transmembrane interface (Figure 5). Thus, residues in or near the proposed state-dependent cholesterol site were implicated in transitions both from and towards the closed state.

**Figure 5.**
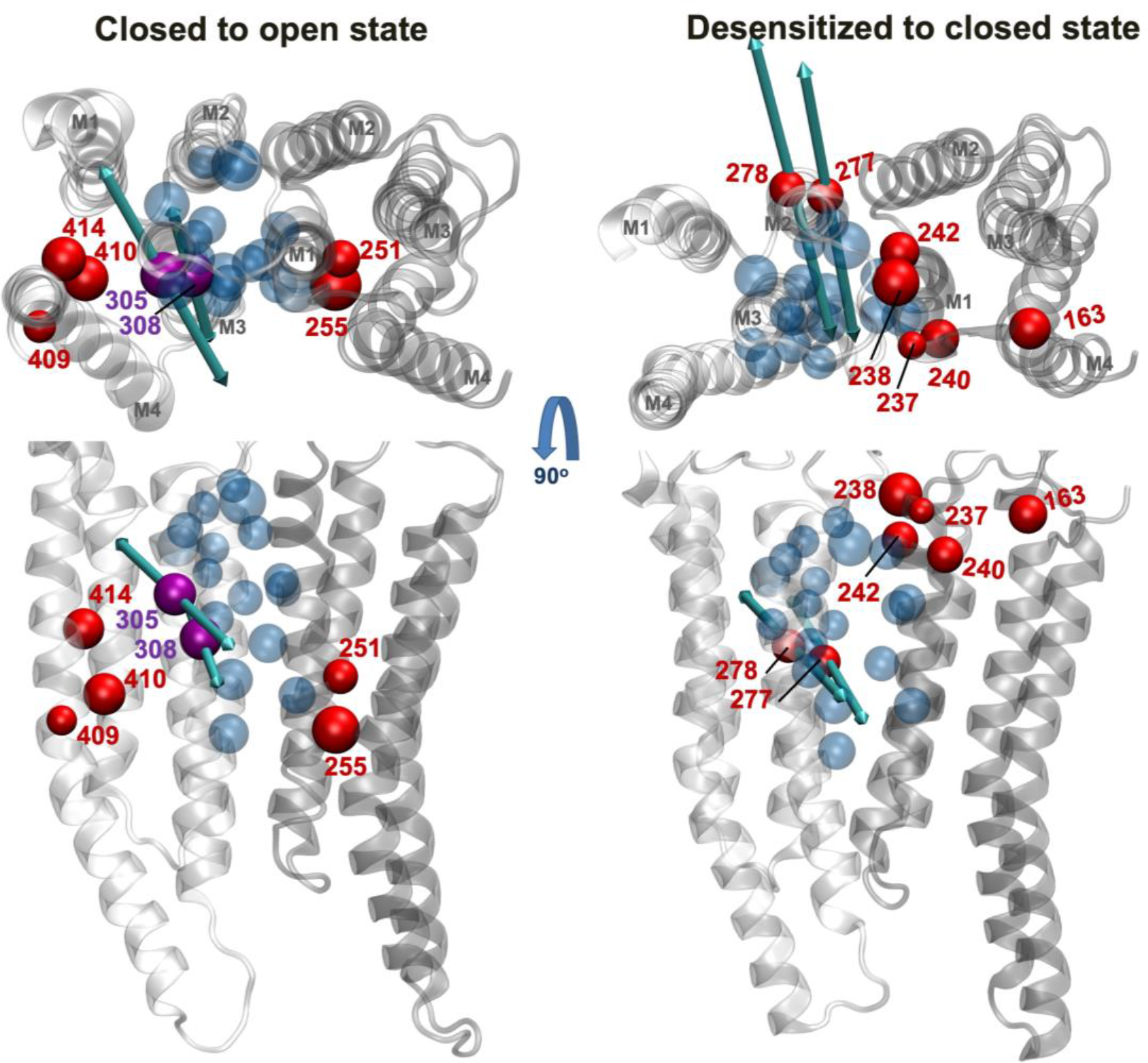
Cholesterol-binding region implicated in channel gating by protein perturbation analysis. A single TMD subunit interface of the a1 GlyR is shown (gray ribbons), with key residues implicated in channel opening (closed-to-open transition, left) and in recovery from desensitization (desensitized-to-closed transition, right) shown as Cα spheres (red). Views are from the extracellular side (top) and from the membrane plane (bottom). Residues Val-305 and Leu-308 (purple) were identified both as key residues in channel opening and as cholesterol contacts. Force vectors corresponding to the two residues with highest overlap are shown as bidirectional arrows, colored cyan. For comparison, residues contacting cholesterol in atomistic simulations (as identified in Figure 4) are shown as transparent blue Cα spheres in VMD bead representation.

### Differential cholesterol binding in heteromeric GlyRs

To test the relevance of our proposed cholesterol interactions in a physiologically potentially more relevant native heteromeric GlyR, we also ran coarse-grained simulations on the pig α1β structure recently reported in a desensitized state (PDB ID 7MLY).^13^ In this system, cholesterol molecules frequently occupied outer-leaflet sites at α-α and α-β interfaces (Figure 6a) overlapping those observed in homomeric receptors, proximal to α1 Ser-283. Cholesterol interactions at these sites were also among the longest in duration (Figure 6b). Interestingly, cholesterol exhibited differential behavior at the β-α interface, where the equivalent residue to Ser-283 is cysteine. In this region, a site with cholesterol present for more than 3 μs duration (BS5) was still observed, but located superficially on the surface of the β subunit rather than buried at the subunit interface.

**Figure 6.**
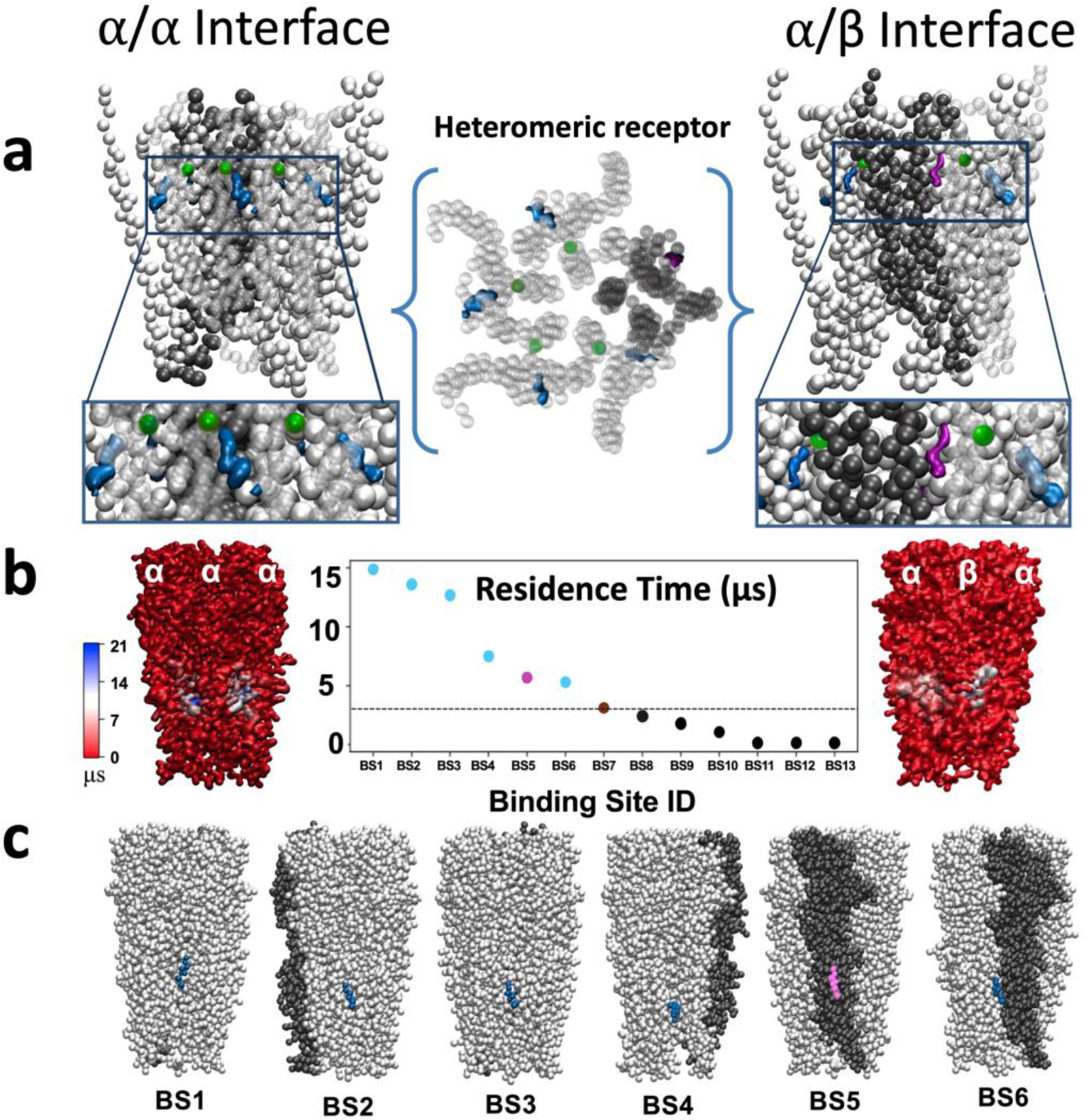
Differential cholesterol binding in heteromeric GlyRs. a) Densities representing >50% cholesterol occupancy around α1 (blue) and β (purple) subunits of a heteromeric GlyR (gray), including a sphere representing Ser- 283 (green). Center view is from the extracellular side; left- and righthand views are from the membrane plane, focusing on different interface types. Insets show zoom views of the outer-leaflet interfacial site. **b)** Interaction residence times of cholesterol at the top thirteen PyLipID binding poses (center), with molecular surfaces focusing on different interfaces at left and right, colored as in Figure 1c. Interaction times are shown for intersubunit sites near the principal face of α1 (blue) or β (purple) subunits, as well as alternative sites of more (brown) or less (gray) than 3 μs (threshold line). Poses with the six longest residence times (BS1–BS6) are illustrated below, showing representative states of the protein (gray) and cholesterol near the principal face of α1 (blue) or β (purple) subunits. One α-α interface is populated by two discrete cholesterol poses (BS1, BS3).

## Discussion

Allosteric gating and modulation are critical to the function and pharmacology of many membrane proteins, including pLGICs. Although 3D structures are increasingly available for this receptor family, intrinsically dynamic allosteric processes may not be clearly described by static structural data, in particular not in the context of interactions with other molecules such as lipids. By combining coarse-grained and atomistic simulations as well as perturbation-based computational analyses of three structures determined under identical experimental conditions, we defined molecular details and mechanistic relevance of a state-dependent cholesterol binding site in a zebrafish GlyR. Additionally, we report that the pig heteromeric GlyR, with 99% identity to the human protein sequence, contains a similar putative binding site at all subunit interfaces except β-α.

In our simulations, cholesterol was capable of intercalating into the outer transmembrane subunit cleft, deep enough to interact directly with residues from the principal-face pore-lining M2 helix. Residue Ser-283 (15’, Ser-267 in human) emerged as a particularly informative location for state-dependent binding, often capable of forming a hydrogen bond with the cholesterol hydroxyl group in simulations of open or desensitized receptors. Differential orientation of this residue in the closed state appeared to preclude cholesterol interactions, reducing cholesterol occupancy in the interfacial site. Sequence variation at this position in GlyR β subunits was further associated with more superficial interactions with this component of a heteromeric receptor. Mutations at Ser-283 have long been shown to alter glycine sensitivity, ^51^ and are linked to autosomal dominant hereditary hyperekplexia, a rare but potentially lethal neuromotor disorder caused by abnormal glycinergic transmission ^52^ (Figure 7a–b). Cholesterol also made frequent polar contacts with the M2 residue Arg- 287 (19’, Arg-271 in human), one turn outward from Ser-283 at the extracellular mouth of the pore. This residue has been termed a “gating mutation,” as mutations appear to uncouple agonist binding from gating, reduce agonist efficacy, and even convert partial agonists into antagonists. Arg-287 is also one of the most common sites of hyperekplexia mutations ^52,53^ (Figure 7a–b).

**Figure 7.**
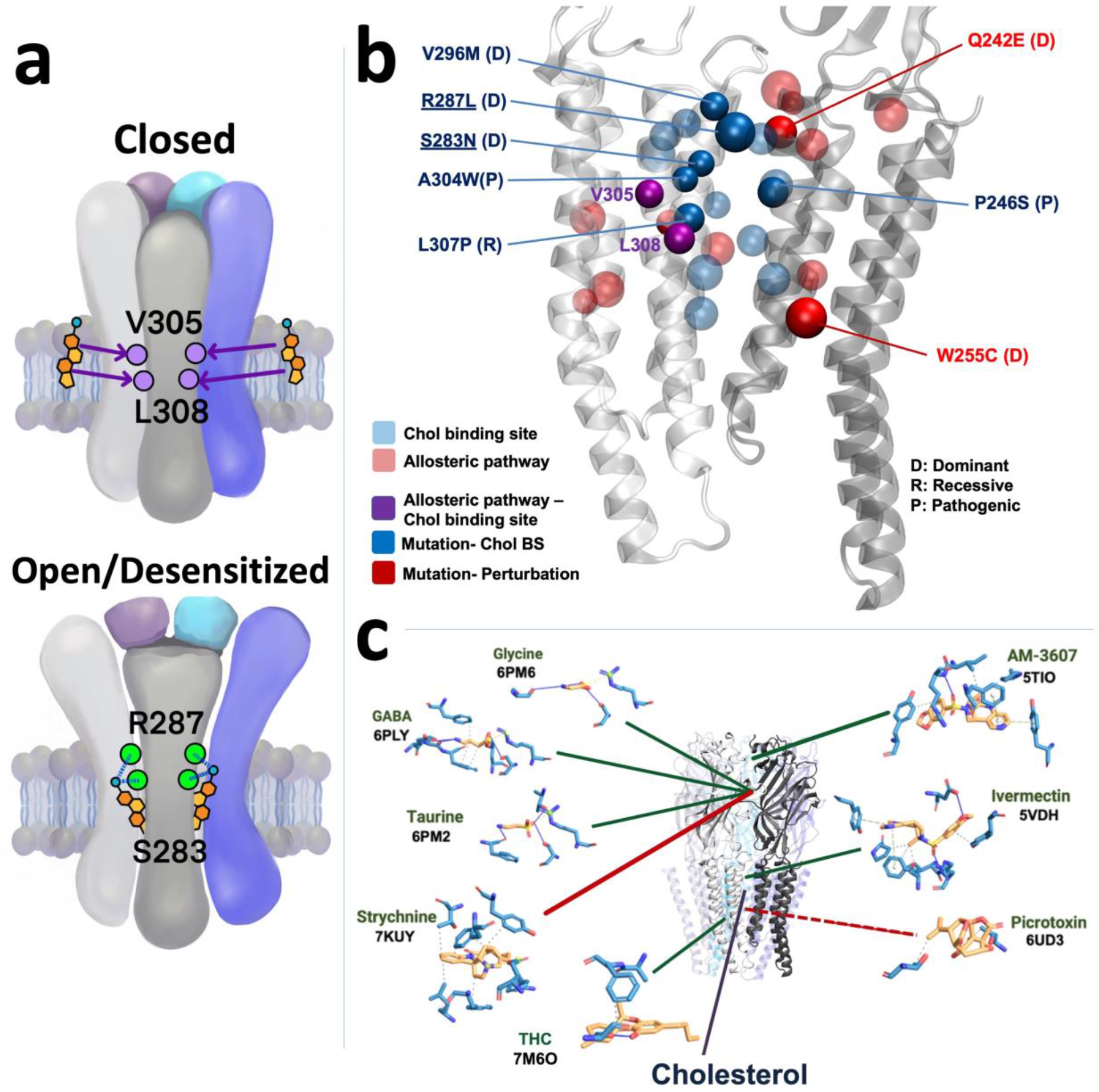
Functionally important GlyR residues implicated in cholesterol binding and/or allostery. a) Proposed state-dependent site for cholesterol, with a preference for binding in the open and desensitized states over the closed state. The top model indicates the potential involvement of residues such as Val-305 and Leu-308 (purple) in binding cholesterol (based on atomistic simulations) and driving channel opening (based on perturbation analysis). The bottom model indicates potential stabilizing contacts of cholesterol in the open and desensitized states with polar residues Ser-283 and Arg-287 (green) on the pore-lining M2 helix. **b)** Solid spheres indicate correspondence between previous evidence for functional relevance and residues implicated here in cholesterol binding (blue), allosteric transitions (red), or both (purple). Residues linked to hyperekplexia are labeled as dominant (D), recessive (R), or indeterminately pathogenic (P). The right-hand labels are associated with M1 of the complementary face, while the left-hand ones correspond to M2–M3 of the principal face; underlined residues are located in M2. **c)** Ligands (gold) resolved in past experimental GlyR structures, with directly coordinating residues (blue). For clarity, the two proximal subunits are shown in gray and black; remaining are semi-transparent.

Extending from the channel pore towards the membrane, several residues making hydrophobic contacts with cholesterol in simulations have also been implicated in channel function (Figure 7a). On the principal M3 helix, substitutions at Val-296 (to Met) and Ala-304 (to Trp) produce tonically open leaky channels, while those at Leu-307 alter glycine sensitivity, surface expression, and/or desensitization. ^54–56^ Mutations at Val-296 and Leu-307 are also associated with hyperekplexia. ^57,58^ On the complementary M1 helix, a hyperekplexia-linked Ser substitution at Pro-246 decreases glycine sensitivity and surface expression. ^54^ The substitution P246S, heterozygous with R81W, also produces fast-desensitizing receptors. ^52,54^ Similarly, substituting a bulky Trp one turn inward at Leu- 249 enhances desensitization in the presence of ivermectin. ^55^ Thus, although functional evidence for cholesterol contacts in GlyRs is limited, specific state-dependent interactions identified in this work are consistent with an influential region for channel function. Peripheral to these direct cholesterol contacts, several of the residues (Gln-242, Tyr-244, Trp-255, Ala-267) we identify (Table 3) as relevant for allosteric gating have also been linked to hyperekplexia ^52,54,58–61^ (Figure 7b). Substituting Glu at M1 residue Gln-242 produces leaky channels, possibly due to enhanced interaction with M2 residue Arg-287 described above. ^54,58^ Conversely, substituting Glu at M2 residue Ala-267 in the inner mouth of the pore suppresses recovery from desensitization. ^61^ These effects support the utility of perturbation analysis in identifying amino acid residues critical to channel function, independent of cholesterol.

The outer transmembrane site we associated with cholesterol may overlap sites for other modulators (Figure 7c). In particular, structures of the GlyR bound to the lipophilic potentiator ivermectin reveal direct interactions with Ile-241, Gln-242, Pro-246, Ser-283, and Ala-304, all frequent contacts of cholesterol. Indeed, residues Pro-246 and Ala-304 were shown to be crucial determinants of ivermectin sensitivity in a TMD mutagenesis screen; the extreme substitution A304F eliminates ivermectin activity altogether ^55,62^. Smaller drugs such as alcohols, anesthetics, cannabidiol, and quercetin have also been associated with the same region in various pLGICs, including influential effects of mutations at Thr-280, ^56,63^ Ser-283, ^3^ Met-303, ^63^ and Ala-304. ^62,64^ Di-cysteine cross-linking studies show that Ala-304 is proximal to both Ser-283 and Ile-245, with the I245C/A304C mutants display unusual responses and S283C/A304C mutation producing leaky channels which function normally after reduction of the disulfide bond These functional results support a role for the outward-facing subunit interface in GlyR gating and drug modulation, particularly residues on M2 and M3 that are also implicated here in cholesterol binding ^65^.

Tetrahydrocannabinol (THC) was also recently observed in a GlyR transmembrane site in contact with residue Phe-410, ^66^ which was implicated here in GlyR opening. Interestingly, cholesterol depletion has been shown to mimic disruption of THC binding. ^3,67^ It is plausible that this drug modulates channel function in part by piggy-backing a site evolved for state-dependent binding of endogenous cholesterol. Recent coarse-grained simulations (in the absence of cholesterol) support a similar intrasubunit site for THC binding and indicate the endogenous cannabinoid N-arachidonyl- ethanolamide (AEA) binds in both this THC site and the intersubunit ivermectin site. ^64^

Our simulations offer testable hypotheses for cholesterol modulation of GlyRs, an effect for which evidence has been limited. In contrast to proteins such as dopamine transporters, GPCRs, and sodium–potassium pumps ^68–70^ with which cholesterol has been shown to crystallize, cholesterol binding to GlyRs has yet to be definitively observed in experiments. Lipidic densities in GlyR structures have generally been modelled as partial phospholipids. ^11,71^ Interestingly, the intersubunit site identified in our work does not contain the proposed cholesterol-sensing CRAC or CARC motifs. ^72,73^ Experiments using mass spectrometry identified cholesterol contacts in the pre-M1, M2–M3 loop, intracellular, and M4 regions, but not in the intersubunit cleft identified here; given that the relevant protocol was likely limited to closed receptors, it is plausible that alternative interactions in that state poise cholesterol for entering the intersubunit pocket upon activation—although simulations indicated such sites are only weakly occupied, even in closed channels. ^74^

Our findings build on those from previous coarse-grained simulations, particularly from homology models of closed and open human GlyRs. ^28^ Whereas that work offered a detailed methodological pipeline and characterized interactions with multiple lipids, here we proceeded from coarse-grained observations to detail amino-acid contacts and dynamics of allostery using atomistic simulations and perturbation analysis. Moreover, since the initiation of that work several new GlyR experimental structures have emerged, in some cases casting doubt on the physiological relevance of homology-modeling templates. Our present work is based on three structures determined under identical conditions, allowing insights into the desensitized state as well as a potentially more relevant open state. In future studies, it may be particularly interesting to probe functional consequences of residues such as Val-305 and Leu-308, implicated here both in binding cholesterol and driving channel opening (Figure 7a).

These findings could further support the discovery of new ligands that are able to modify or substitute for cholesterol interactions. Various hydrophobic compounds including coenzyme Q10, α- tocopherol, vitamins D3 and K1 can attach to cholesterol sites. ^17^ In particular, GlyRs are modulated by several steroid hormones, which are structurally similar to cholesterol and could function through shared binding sites. Stress hormones including corticosterone, cortisol, and their metabolites have been shown to promote GlyR desensitization in neurons, potentially linking to brain dysfunction in chronic stress. ^75^ Moreover, modulation of GlyRs by neurosteroids appears to be subunit-dependent: the presence of β subunits reduces their sensitivity, possibly due to a decrease in available binding sites ^76^ as indicated here for cholesterol.

## Data Availability

Simulation frames and parameters for both coarse-grained and atomistic simulations can be accessed via the DOI: 10.5281/zenodo.8374103.

## Acknowledgments

This work was supported by grants from the Swedish Research Council (2019-02433, 2021-05806), the BioExcel Centre of Excellence for Computational Biomolecular Research (EuroHPC Joint Undertaking 101093290), the Swedish e-Science Research Center, and compute resources from the National Academic Infrastructure for Supercomputing in Sweden (SNIC 2022/3-40, NAISS 2023/3- 27).

