## Supplemental Data 1 for "An Allosteric Cholesterol Site in Glycine Receptors Characterized Through Molecular Simulations"

### Supporting Information

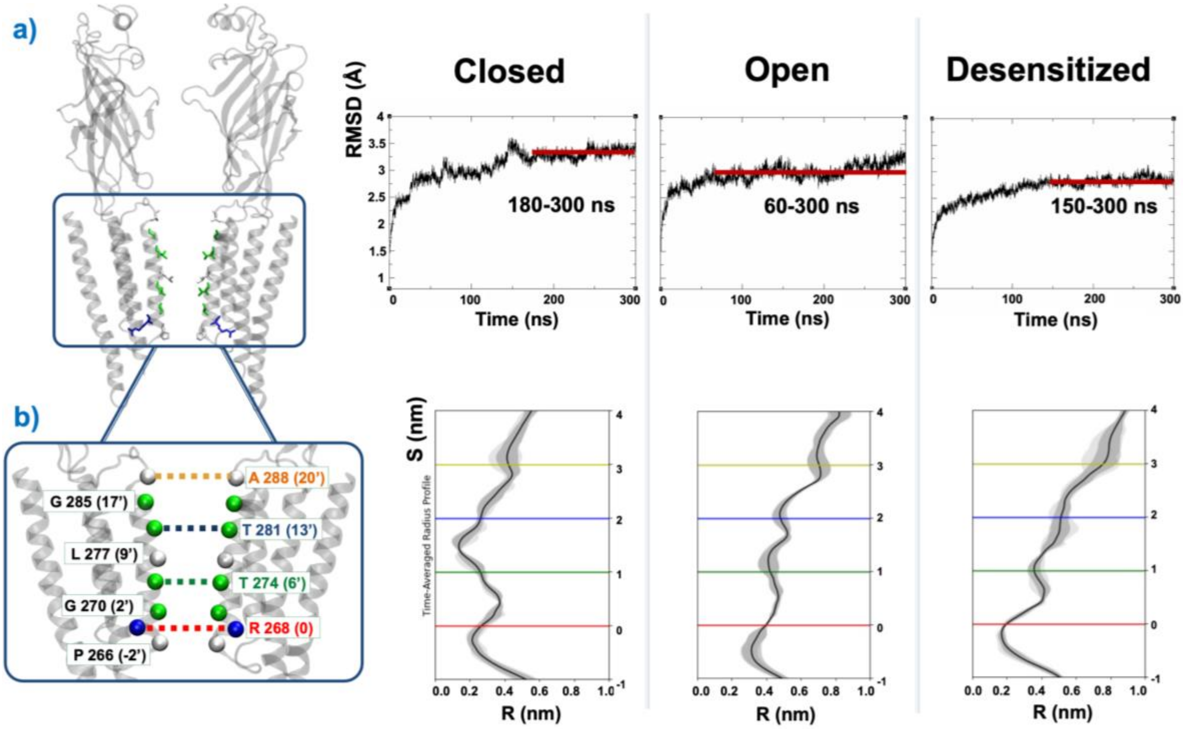


**Figure S1. Structural stability of GlyR models evaluated through RMSD analysis and pore characterization. a)** The RMSD of closed (PDB ID 6PM3), open (PDB ID 6PM2), and desensitized (PDB ID 6PM1) receptor states was measured to assess the structural stability of the entire protein. The RMSD values are plotted over time and the equilibrated chunk of the simulation, indicated by the red line, was selected for subsequent perturbation analysis (right). Two facing chains of protein are illustrated in a gray cartoon representation, while the pore lining residues are depicted in licorice, and color-coded according to their residue types using VMD. **b)** The pore radius along the permeation pathway is measured to ensure the pore stability throughout the simulation using CHAP. The arc length along the central axis of the conduction pathway is plotted with error bars, with residue R288 defined as the reference point at zero (right). For clarity, the regions at 0, 6', 13', 20' within the pore in the zoomed-in view (left) are color-coded using the same color scheme: red, green, blue, and yellow, respectively.

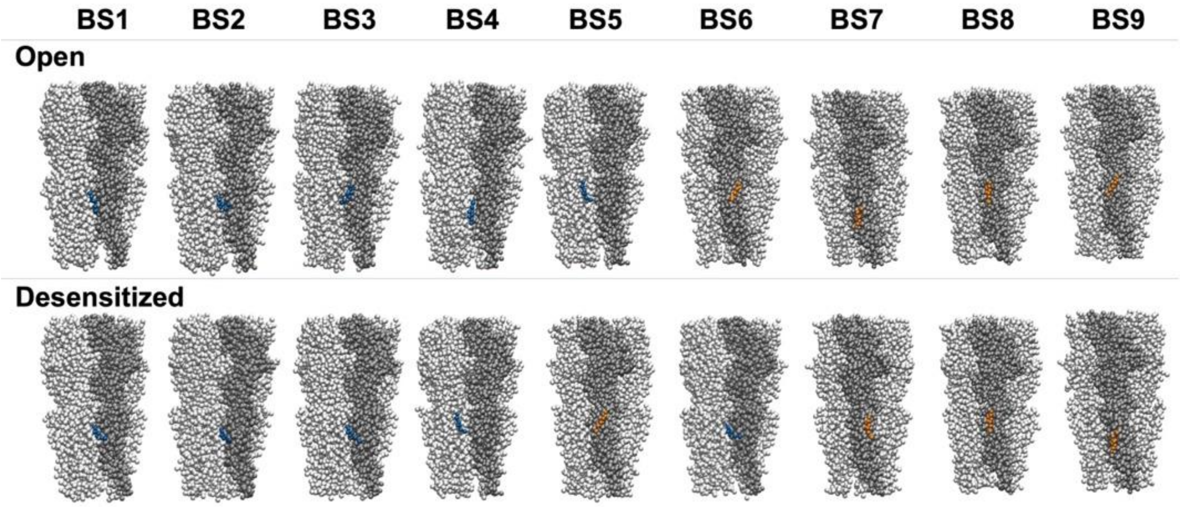


**Figure S2. PyLipID analysis revealed nine distinct cholesterol interaction poses across the two states.** Coarse-grained simulations were conducted using PyLipID to uncover the optimal cholesterol binding poses (BS1-BS9) with the open (top) and desensitized (bottom) states of the α1 GlyR, employing density-based scoring and clustering methods. This approach unveiled nine distinct poses, with those located within the intersubunit region highlighted in blue and those in other regions displayed in yellow. Poses with interaction times shorter than a 3 μs threshold were excluded from consideration as binding events and are not displayed.

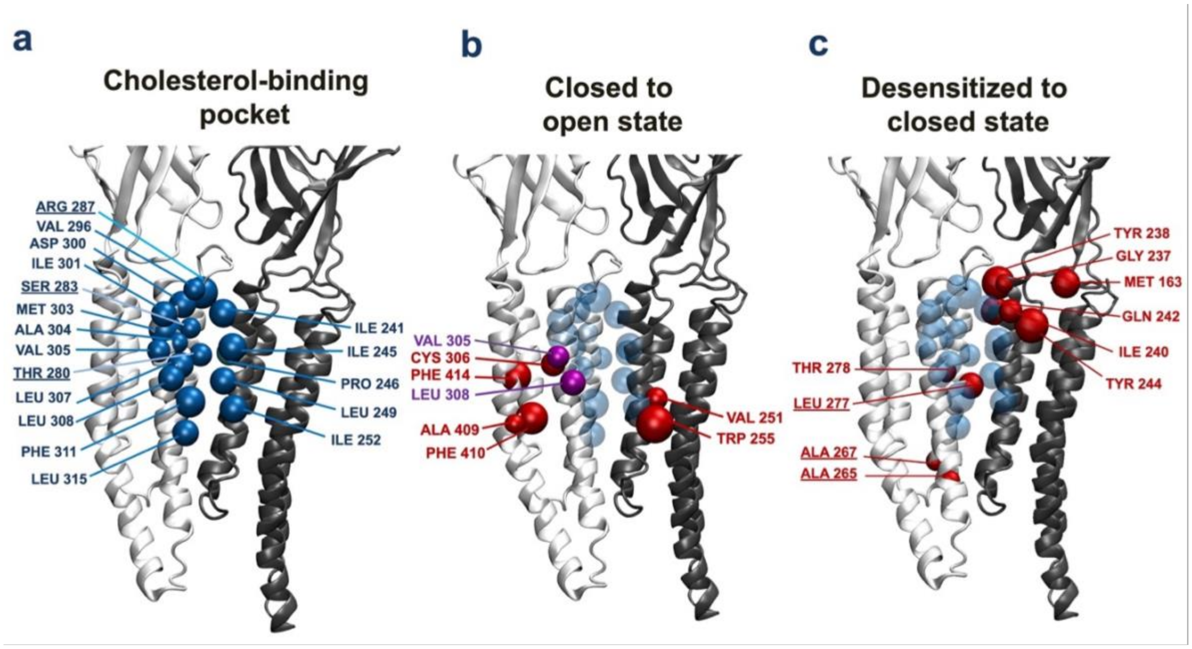


**Figure S3. Potentially allosterically relevant residues. a)** Coarse-grained simulations were utilized to pinpoint regions involved in cholesterol binding. These were refined through back-mapping followed by atomistic simulations to provide insight into specific residues, visualized as blue spheres, contributing to the cholesterol binding site. **b)** Residues correlated with the closed-to-open state transition in the perturbation analysis are underlined. Residues that overlap with the cholesterol binding site (as depicted in *a* are represented as purple spheres, while the remaining ones are colored in red. **c)** Residues correlated with the desensitized-to-closed state transition are highlighted as red spheres in VMD bead representation. For clarity, residues at the cholesterol binding site are depicted as transparent blue spheres. Two adjacent chains are depicted in cartoon representation and colored in black and white using VMD.
